# A systems vaccinology resource to develop and test computational models of immunity

**DOI:** 10.1101/2023.08.28.555193

**Authors:** Pramod Shinde, Ferran Soldevila, Joaquin Reyna, Minori Aoki, Mikkel Rasmussen, Lisa Willemsen, Mari Kojima, Brendan Ha, Jason A Greenbaum, James A Overton, Hector Guzman-Orozco, Somayeh Nili, Shelby Orfield, Jeremy P. Gygi, Ricardo da Silva Antunes, Alessandro Sette, Barry Grant, Lars Rønn Olsen, Anna Konstorum, Leying Guan, Ferhat Ay, Steven H. Kleinstein, Bjoern Peters

**Affiliations:** Center for Infectious Disease and Vaccine Research, La Jolla Institute for Immunology, La Jolla, CA, USA; Bioinformatics and Systems Biology Graduate Program, University of California, San Diego, CA, USA; Department of Health Technology, Technical University of Denmark, Kongens Lyngby, Denmark; Knocean Inc., 107 Quebec Ave. Toronto, Ontario, M6P 2T3, Canada; Program in Computational Biology & Bioinformatics, Yale University, New Haven, CT, USA; Department of Medicine, University of California, San Diego, San Diego, CA, USA; Department of Molecular Biology, School of Biological Sciences, University of California San Diego, La Jolla, California, USA; Department of Pathology, Yale University School of Medicine, New Haven, CT, USA; Department of Biostatistics, Yale School of Public Health, New Haven, CT, USA

## Abstract

Computational models that predict an individual’s response to a vaccine offer the potential for mechanistic insights and personalized vaccination strategies. These models are increasingly derived from systems vaccinology studies that generate immune profiles from human cohorts pre- and post-vaccination. Most of these studies involve relatively small cohorts and profile the response to a single vaccine. The ability to assess the performance of the resulting models would be improved by comparing their performance on independent datasets, as has been done with great success in other areas of biology such as protein structure predictions. To transfer this approach to system vaccinology studies, we established a prototype platform that focuses on the evaluation of Computational Models of Immunity to Pertussis Booster vaccinations (CMI-PB). A community resource, CMI-PB generates experimental data for the explicit purpose of model evaluation, which is performed through a series of annual data releases and associated contests. We here report on our experience with the first such ‘dry run’ for a contest where the goal was to predict individual immune responses based on pre-vaccination multi-omic profiles. Over 30 models adopted from the literature were tested, but only one was predictive, and was based on age alone. The performance of new models built using CMI-PB training data was much better, but varied significantly based on the choice of pre-vaccination features used and the model building strategy. This suggests that previously published models developed for other vaccines do not generalize well to Pertussis Booster vaccination. Overall, these results reinforced the need for comparative analysis across models and datasets that CMI-PB aims to achieve. We are seeking wider community engagement for our first public prediction contest, which will open in early 2024.

## Introduction

A common challenge in developing computational models for biological applications is to objectively test their generalizability and predictive performance^1–3^. This is especially true for systems vaccinology studies, due to the heterogeneous and high-dimensional nature of the assay readouts, differences in study designs, and incomplete understanding of what the clinically relevant correlates of a vaccine induced protective response are. Integrating diverse data types, accounting for inter-individual variability, and capturing temporal dynamics are crucial aspects that need to be addressed to ensure the robustness and accuracy of computational models in system vaccinology.

To address these challenges, we have established the CMI-PB resource to develop and test computational models that predict the outcome of Tdap booster vaccination that is designed to be used by the broader community. As part of this project, we will measure the response to Tdap booster vaccination over four years, in four independent donor cohorts datasets, which will allow us to create and test computational models that predict vaccination outcome based on the baseline state of the vaccines (Table 1).

**Table 1:** Past and future CMI-PB annual prediction challenges. Our commitment involves conducting four annual challenges. The first challenge was completed in May 2022 with participation from the CMI-PB consortium. The second challenge will be announced in August 2023 and will feature the CMI-PB consortium along with a limited number of invited contestants from outside the consortium. We will involve members of the public in the third and fourth challenges. The first challenge included training data from a previously published study^14^ and newly generated test data. Similarly, we will use both the training and test data from previous challenges as the training data for future challenges and generate new data for testing purposes.

|  | Annual prediction challenge title | Contestants | Number of subjects |  | Current status |
| --- | --- | --- | --- | --- | --- |
|  |  |  | Training dataset | Test dataset |  |
| 1 | **First Challenge:**<br>Internal dry run | CMI-PB consortium | 60 (28 aP + 32 wP) | 36 (19 aP + 17 wP) | Concluded in May 2022 |
| 2 | **Second Challenge:**<br>Invited challenge | Invited contestants | 96 (47 aP + 49 wP) | 23 (13 aP + 10 wP) | Will be announced in September 2023 |
| 3 | **Third Challenge:** | Public | 119 (60 aP | 32 (16 aP + | Will be announced in April |
|  | Open Challenge 1 |  | + 59 wP) | 16 wP) | 2024 |
| 4 | **Fourth Challenge:**<br>Open Challenge 2 | Public | 151 (76 aP<br>+ 85 wP) | 32 (16 aP +<br>16 wP)* | Will be announced in<br>December 2024 |

\*Goal

Here we report on the outcome of the first challenge: an internal ‘dry run’ where the teams involved in making predictions spanned multiple institutions, but were all collaborators in developing the CMI-PB resource. We report on the challenges encountered for data sharing, formulating prediction questions, and the interpretation of the results from different prediction models including the determination of which factors contributed to such predictions. These results will inform the design of the next prediction contest, which will open to community participation at the end of 2023.

The overall goal of our study is two-fold: First, we want to establish a community platform to test and compare computational models of immunity in vaccination. Second, we want to better understand vaccine induced immunity to *B. pertussis* - the causative agent of whooping cough - a highly contagious respiratory infection that causes severe disease in infants^4^. In the past 3 decades, there has been a resurgence of whooping cough infections in the US that has been linked to a switch in the vaccine from wP (whole cell) to aP (acellular) in 1996. Epidemiologically, that suggested that the immunity induced by aP vaccines wanes quicker than those by wP vaccines^5–9^. A number of differences in immune responses between aP and wP vaccines have been identified, such is their T cell polarization^10–13^, but it is unclear how those differences translate to duration of immunity. By establishing and testing computational models that attempt to predict the cascade of events that follow *B. pertussis* booster vaccination, we will improve our understanding of the mechanisms underlying these events, with the ultimate goal to identify what variables induce a strong and durable recall response.

## Results

This results section covers two components: First, we describe the experience in setting up and running the prediction contest. Second, we describe specific models that were developed and discuss their performance on the prediction tasks.

### Experimental data generation and access

Our experimental study is designed for a systems-level understanding of the immune responses induced by Tdap booster vaccination and closely mimics the design of previous studies from our group^14^. Briefly, individuals primed with aP or wP in infancy-were boosted with Tdap and blood was collected pre-booster and post booster at days 1, 3, 7, and 14 (**Figure 1A**). Multiple assays were performed, namely i) gene expression analysis (RNAseq) of bulk peripheral blood mononuclear cells (PBMCs), ii) plasma cytokine concentration analysis, iii) cell frequency analysis of PBMC subsets, and iv) plasma antibodies against Tdap components. We previously generated data from a total of 60 subjects (28 aP and 32 wP; **Table 1**) as part of ^14^ that was made available as a training dataset to develop predictive models. Additionally, we generated data from a separate group of 36 newly tested subjects (19 aP + 17 wP) that was set aside to serve as test data for the predictions.

**Figure 1:**
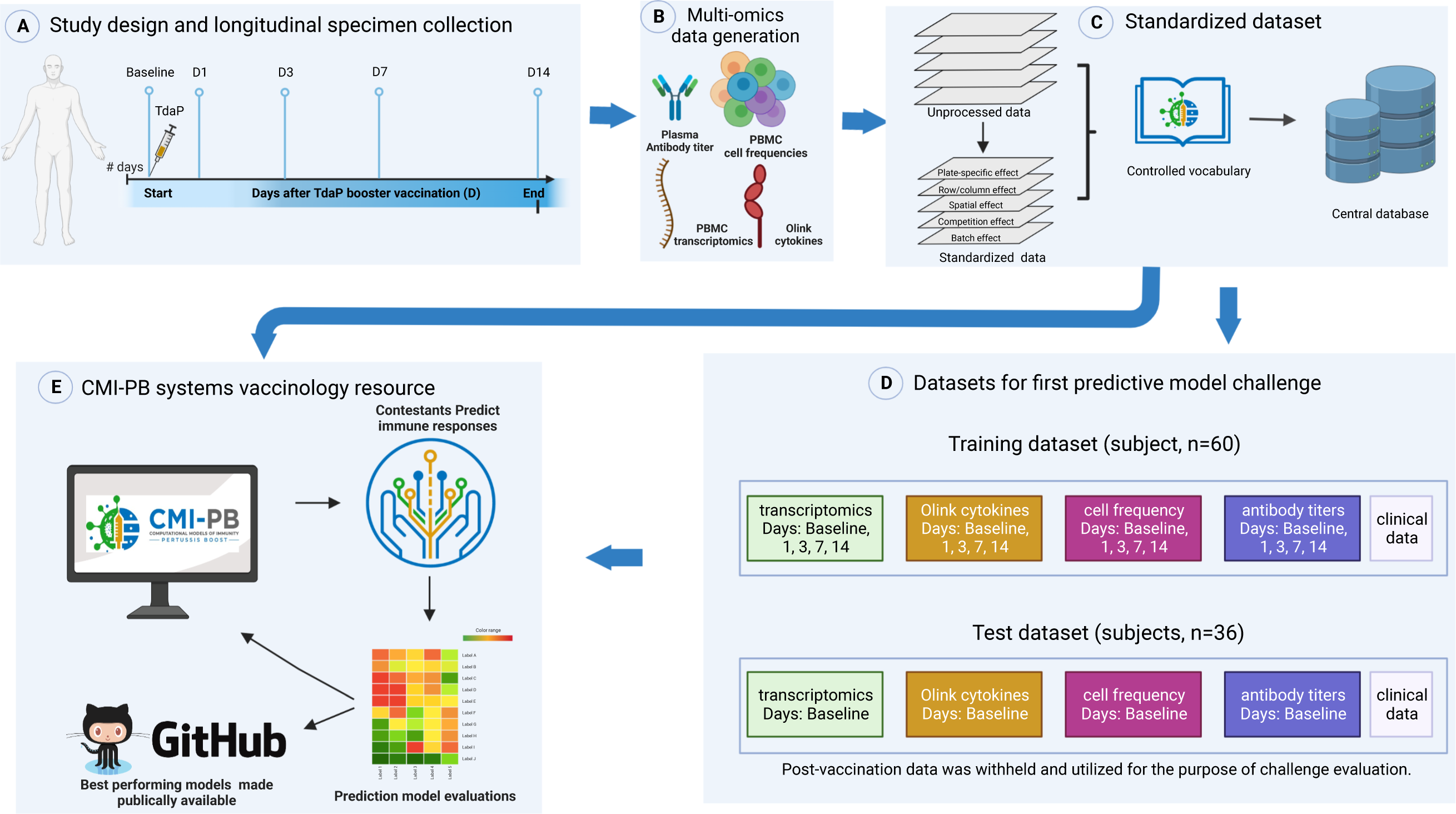
Outline for establishing the CMI-PB resource. A) Recruitment of human subjects and longitudinal specimen collection. B) Generation of multi-omics data to obtain a comprehensive understanding of the collected specimens. C) Implementation of a data standardization approach to ensure consistency and comparability of the generated data. D) The resulting dataset is provided in training and test formats to enable contestants to develop their predictive models. E) The CMI-PB resource website serves as a platform for hosting an annual prediction challenge, offering data visualization tools for generated data, and providing access to teaching materials and datasets.

To integrate experimental data generated at different time points, we created a centralized CMI-PB database with tables corresponding to entity categories, including subject and specimen information, experimental data, and ontology tables (database schema is provided in **Figure S1**). We established different access modalities including an application programming interface (API; https://www.cmi-pb.org/docs/api/) and bulk file downloads, and shared these different access modalities with our participants in the prediction challenge.

### Formulating the prediction tasks

We formulated multiple prediction tasks in order to quantitatively evaluate different aspects of immune responses to Tdap booster vaccination. As targets, we selected biological readouts known to be changed by booster vaccination under the premise that they are likely to capture meaningful heterogeneity across study participants based on our previous work^14^. For instance, we had previously shown that the percentage of monocytes was significantly elevated on day 1 post-booster vaccination compared to baseline (i.e., pre-booster vaccination)^14^. We created a first task in which the overall frequency of monocytes among PMBCs on day 1 post-booster vaccination has to be predicted. Similarly, we had previously shown that plasma IgG1-4 levels significantly increased at day 7 post-booster vaccination compared to baseline^14^, and accordingly chose as another task to predict plasma IgG levels against the Pertussis toxin (PT) on day 14 post-booster vaccination. The third task was based on our previous finding that a subset of aP-primed individuals showed an increased expression of pro-inflammatory genes, including *CCL3* on day 3 post-booster vaccination^14^, which was the target of the third task. Overall, we formulated a total of 14 prediction tasks (**Table S1**), including 13 prediction tasks of readouts identified from previous work and a “sanity-check” task to predict the expression of the sex-specific *XIST* gene post-booster vaccination per individual^15^.

### Choosing a metric to evaluate prediction performance

We set out to choose a <u>metric</u> to evaluate how different prediction methods performed. Specifically, we wanted to have three considerations: i) we needed a metric that would produce a single numeric value as an output. This would allow us to compare and rank the performance of the prediction methods effectively, ii) the chosen metric needed to be non-parametric because the different experimental assays utilized in the study produce analyte measurement outputs with non-normal distributions, iii) we wanted to avoid incorporating arbitrary cutoffs or thresholds that could introduce subjectivity or bias into the assessment process. Based on these considerations, we chose the Spearman Rank correlation coefficient as our primary metric. The prediction tasks in our first challenge thus constituted predicting the rank of individuals in specific immune response readouts from high to low after *B. pertussis* booster vaccination based on their pre-vaccination immune status.

### Feedback from participants prior to data submission

We shared the prediction tasks, metrics, and data access instructions with the contest participants, and asked them for feedback prior to submitting their prediction results. Two main points of feedback were: (1) All contestants preferred using the bulk file downloads over utilizing the custom API we had created, as they preferred to work with raw data over having to learn a new access modality. Given that creating reliable APIs is resource intensive, this was identified as an area we wanted to down-prioritize going forward. (2) When inspecting antibody titer data generated in different years, contestants noticed significant variation in the averages of the baseline values for donors (subjects) between the test and training datasets. Those variations were due to a switch in the site where the assays were performed. We thus standardized the antibody data in each year by applying the baseline median as a normalization factor (https://github.com/CMI-PB/2021-Ab-titer-data-normalisation; **Figure S2, S3**), and provided both the raw data and normalized data to the contestants.

### Gathering and evaluating prediction results

A total of 34 computational models were developed by three independent teams of contestants. Each team worked separately on their own set of models. The first team focused on identifying and constructing prediction models based on the systems vaccinology literature (**Figure 2B**). The second and third teams, on the other hand, focused on constructing new prediction models derived from multi-omics dimension reduction techniques (**Figure 2C-D**). We established a deadline of 3 months for each team to submit their models, and subsequently, the corresponding predictions were received for evaluation. A complete submission file contained 14 columns, 1 column per prediction task. We found that most prediction models focused on a subset of tasks. Furthermore, we found that in some cases, predictions for individual donors were omitted. In those cases, we used the median rank calculated from the ranked list submitted by the contestant to fill in missing ranks. An overview of the prediction results is summarized in **Figure 3**, and the different prediction approaches are described in more detail in the following.

**Figure 2.**
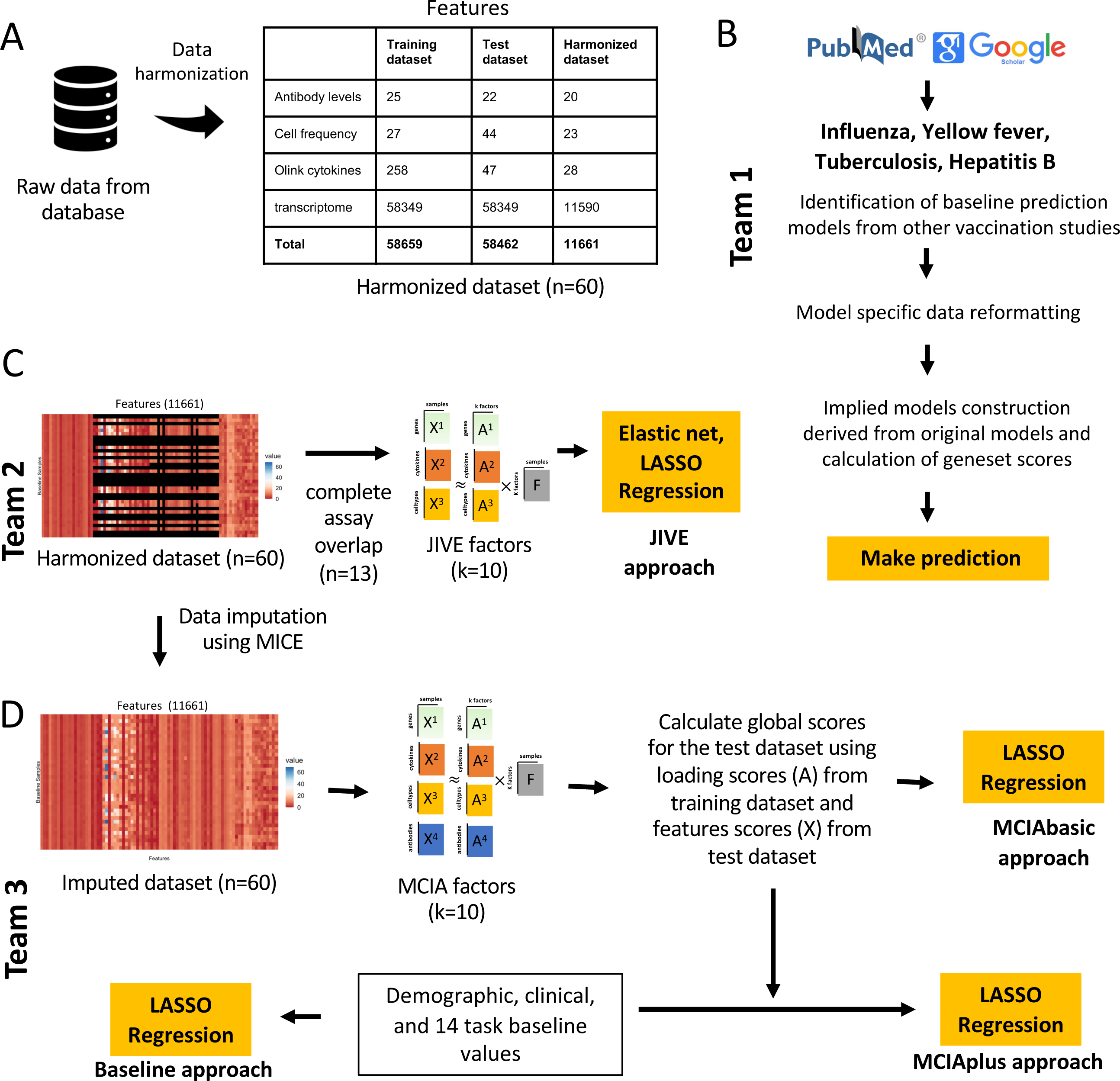
Data processing, computable matrices, and prediction model generation. A) Generation of a harmonized dataset involved identifying shared features between the training and test datasets and filtering out low-information features. Literature-based models used raw data from the database and applied data formatting methods specified by existing models. In contrast, JIVE and MCIA utilized harmonized datasets for constructing their models. B) Flowchart illustrates the steps involved in identifying baseline prediction models from the literature, creating a derived model based on the original models’ specifications, and performing predictions as described by the authors. C) The JIVE approach involved creating a subset of the harmonized dataset by including only subjects with data for all four assays. The JIVE algorithm was then applied to calculate 10 factors, which were subsequently used for making predictions. JIVE employed five different regression models for prediction purposes. D) MCIA approach applied MICE imputation on the harmonized dataset and used this data for model construction. MCIA method was applied to the training dataset to construct 10 factors. Then, these 10 factors and feature scores from the test dataset were utilized to construct global scores for the test dataset. LASSO regression was applied to make predictions. MCIAplus model was constructed by including additional features (demographic, clinical features, and 14 task values) as factor scores, and it also utilized LASSO regression to make predictions. D) The MCIA approach utilized MICE imputation on the harmonized dataset for model construction. The MCIA method employed the imputed training dataset to construct 10 factors. These 10 factors, along with feature scores from the test dataset, were used to construct global scores for the test dataset. LASSO regression was applied to make predictions. Additionally, the MCIAplus model incorporated additional features such as demographic, clinical features, and 14 task values as factor scores. Finally, LASSO regression was employed for making predictions.

**Figure 3:**
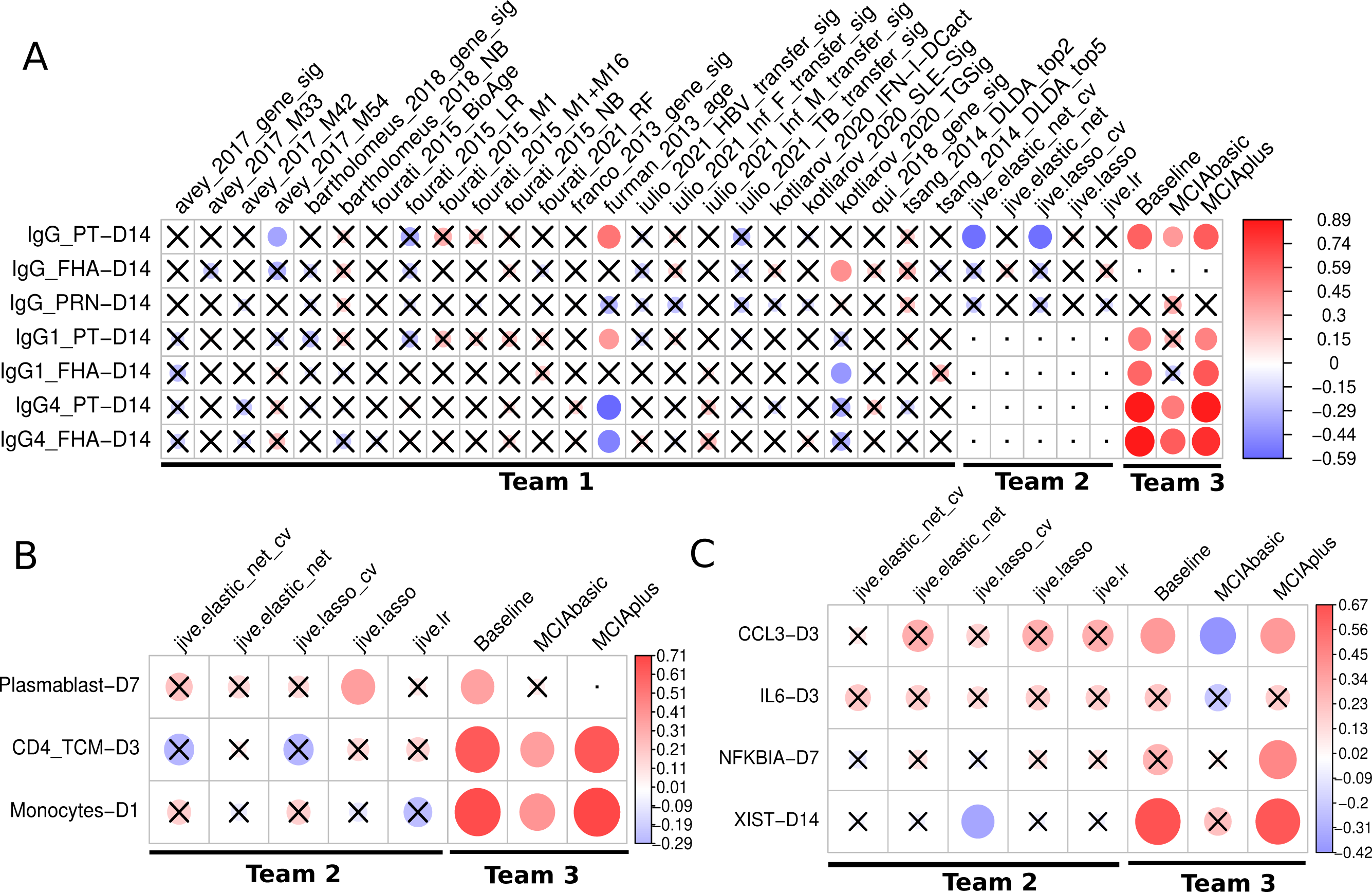
Evaluation of the prediction models submitted for the first CMI-PB challenge. Model evaluation was performed using Spearman’s rank correlation coefficient between predicted ranks by a contestant and actual rank for each A) Antibody titers, B) Immune cell frequencies and C) transcriptomics tasks. The circles in the heatmaps are sized proportionally according to the absolute values of the Spearman rank correlation coefficient, while crosses represent any correlations that are not significant. The dot represents whether the model does not submit ranks for a particular task or if submitted tasks contain unique ranks. Any submissions featuring unique ranks are not included in the evaluation process. The baseline and MCIAplus models submitted by team 3 outperformed other models for most tasks.

### Team 1: Establishing prediction models from the systems vaccinology literature

The first team set out to identify previously published models developed within the systems vaccinology field that aim to predict vaccination outcomes. With systematic keyword queries using PubMed and Google Scholar and following citations, we identified 40 studies of potential interest. A detailed review of these papers identified 10 studies with 24 models that were suitable for our purpose as they i) used pre-vaccination measurements that we have available in our CMI-PB study, ii) established biological differences in vaccine responses that we could utilize in our prediction tasks, which in practice meant predicting antibody titers levels or classifying subjects into high or low vaccine-induced antibody responders ^16–25^. None of the identified models were developed for *B. pertussis*, but rather covered a wide range of vaccines, including those against influenza, hepatitis B, and the yellow fever virus. They employed a variety of methodologies including classification-based (diagonal linear discriminant analysis, logistic regression, Naive Bayes, Random Forest), regression-based (elastic net), and other approaches (gene signature and module scores). A summary of the literature review is depicted in **Figure 2B and Figure S4** for the 24 prediction methods that were implemented (**Table S2**). For each literature model, we adapted the output scores to our prediction tasks, as described in the methods. It has to be re-emphasized that these models were not developed to predict *B. pertussis* vaccination outcomes. Thus evaluating these models as part of our prediction contest does not quantify their intended performance, but rather judges how universal previously identified patterns that impact responses for other vaccines are.

### Establishing a harmonized dataset to train machine learning (ML) models

The second and third team both worked on ML approaches specific to B. pertussis vaccination. While the prediction approaches they chose were different, they collaborated on generating a harmonized dataset to start out with. Specifically, many of the features evaluated in our experimental assays have low-information content. Incorporating less informative features introduces various challenges in data analysis. Low analyte levels could be difficult to distinguish from background noise, missing data could skew statistical analyses, and these features tend to make it more challenging to identify a robust and accurate prediction model. To address these issues, we applied feature filtering on each assay in the training dataset, which is a widely adopted data pre-processing strategy^14^. For gene expression, we filtered zero variance and mitochondrial genes, and removed lowly expressed genes (genes with transcript per million (TPM) < 1 in at least 30% of specimens). Similarly, we filtered features with zero variance from cytokine concentrations, cell frequency, and antibody assays. Subsequently, we removed features not measured for the test dataset and retained only those that overlapped between the training and test datasets. As a result, we were left with a total of 11,661 features in the harmonized dataset out of the original 58,659 features (**Figure 2A**). These harmonized datasets were used for training of the two ML approaches as described below.

### Team 2: Establishing purpose-built models using Joint and Individual Variation Explained (JIVE)

The second team set out to build new prediction models using the available CMI-PB training data using joint dimensionality reduction methods that discover patterns within a single modality and across modalities to reduce the number of dimensions. In particular, we applied the JIVE method to reduce the dimensionality of our datasets before applying regression-based models to make predictions^26,27^. JIVE decomposes a multi-source dataset into three terms: a low-rank approximation capturing joint variation across sources, low-rank approximations for structured variation individual to each source, and residual noise^27^. This decomposition can be considered a generalization of Principal Components Analysis (PCA) for multi-source data^27^. For JIVE, harmonized datasets for transcriptomics, cell frequency, and cytokines concentrations were first intersected on subjects which resulted in 13 individuals with complete data, and finally, the decomposition was applied, generating 10 factors per omics data (**Figure 2C**). These factors were then used as input for five different regression-based methods to turn the JIVE results into predictive models for each specific task. These regression methods included linear regression, lasso and elastic net with default parameters and two more variants of lasso and elastic net that involved an automatic hyperparameter search via cross-validation (CV; see **Figure 2C**).

### Team 3: Establishing purpose-built models using Multiple Co-Inertia Analysis (MCIA)

The third team worked on three different approaches to build prediction models (**Figure 2D**). The first approach (baseline approach) utilized clinical features (age, infancy vaccination, biological sex) and baseline task values as predictors of individual tasks. The second approach (the MCIAbasic) utilized 10 multi-omics factors constructed using MCIA as predictors of individual tasks. Prior to implementing MCIA, the harmonized datasets were further processed to impute missing data in the baseline training set using Multiple Imputation by Chained Equations (MICE) algorithm (**Figure 2**)^28^. The objective function in MCIA maximizes the covariance between each individual omic and a global data matrix consisting of the concatenated omic data blocks^29,30^. Finally, the third approach (MCIAplus) combined the first two approaches and utilized clinical features, baseline task values, utilized baseline approach, and 10 MCIA factors identified through the MCIAbasic approach as predictors of individual tasks. Further, for all three approaches, we built a general linear model with lasso regularization for each task. We used the feature scores as input data and the prediction task values as response variables, generating separate predictive models for each task.

### Comparing model prediction performance

In total, 32 different sets of predictions were submitted, including 24 from Team 1 based on models identified from the literature and 8 models from Team 2 and 3 specifically trained on the available data. A heatmap visualization of Spearman’s correlations for tasks versus models is presented in **Figure 3**. For 10 out of 12 prediction tasks at least one model had a significant correlation of its predictions with the actual results, while no model showed significant correlations for the remaining 2 tasks.

For Team 1, of the twenty-four literature-based models, only three (furman_2013_age, kotliarov_2020_TGSig, and avay_2017_m54) showed a significant correlation for any of the seven antibody-related tasks by at least one model. The most successful model by far was derived from a previous study by Furman et al^20^ (furman_2013_age), where chronological age of an individual was used as the sole predictor for antibody response levels to influenza vaccination. In our results, age correlated positive with IgG1, and negative with IgG4 titers to vaccine antigens post boost. These results suggest that chronological age has a strong predictive potential to be universally utilized as a biomarker to predict antibody responses against different pathogens in addition to influenza and *B. pertussis*.

For Team 2, JIVE-based submissions attempted 10 tasks, excluding the four antibody-related tasks that had missing samples within the harmonized dataset. Diving into the cell frequency tasks, we saw a modest performance for predicting plasmablast levels on day 7, and surprisingly, the simple linear regression performed best. Whereas for other cell frequency tasks, there was no clear pattern of model performance. Within gene expression tasks, JIVE-based models performed best when predicting *CCL3* levels on day 3 and, once again, models without hyperparameter tuning performed the best. Turning to *IL6* at day 3, all JIVE-based models performed modestly, which suggests that this task may be harder than others. As for *NFKBIA* on day 7, predictions were poor for all JIVE-based models. The poor performance of JIVE-based models on some predictive tasks may be due to the limited number of samples (subjects) used (n=13).

For Team 3, the baseline, MCIAbasic, and MCIAplus approaches submitted predictions for all 14 tasks. These approaches outperformed all other teams prediction performance. Specifically, both the MCIAplus and baseline approaches demonstrated significant correlations for 10 out of the 14 tasks, covering the antibody titer, gene expression, and cell frequency tasks - as illustrated in **Figure 3**. The MCIAbasic approach had 6 out of 14 significant correlations. When examining what factors led to improved performance of the baseline and MCIAplus approach as compared to the MCIAbasic approach, it is apparent that the latter does not include demographic and clinical information such as age.

## Discussion

Here, we report the first evaluation of computational prediction models applied to vaccine immune responses to *B. pertussis*. The concept of our approach is based on previous successful efforts such as the CASP and DREAM challenges^31,32^, which have shown that community prediction contests can significantly advance the field of computational predictions. This inaugural “dry run” constitutes an important step for the development and refinement of our future community prediction contests. Major lessons were learned both on how such a contest should be run, and on what input variables should be considered in successful prediction models.

For future contests, we received a clear message that we should focus on simplicity. Contestants asked for fewer prediction tasks, as each task has additional overhead. In examining the results multiple related tasks tended to show similar results, which further confirms that fewer unique tasks are preferable. For data access, there was a clear preference for direct data downloads as files that are familiar to bioinformaticians over the use of novel APIs. We will take this into account. Finally, data harmonization/ normalization is a key concern. Even in a tightly controlled project like ours, we cannot guarantee that the data we generated in year one can be generated the same way in year two. Initially, we did not expect this to be a problem, but instruments eventually break, reagents run out, and the companies supplying them can go bankrupt or change what they make available. This makes it crucial for data generating sites to document how data was generated and to provide a way to harmonize it with previous data.

For model building, we were intrigued that of all the models assembled from the literature, the best prediction performance was based solely on chronological age. Aging is characterized by a progressive loss of physiological integrity and an increased susceptibility to immunosenescence^33^. Age has been reported to be an important determinant of vaccine effectiveness in older adults^34^. So it is no surprise that age is associated with vaccine efficacy, but it is surprising that none of the more comprehensive models that take into account other variables - such as inflammation - perform better, which we would have expected to be the case, even if these other models were developed for different vaccines.

For the two ML approaches implemented by our teams, JIVE and MCIA ML, the most striking difference in performance was clearly due to how baseline values were included in the MCIA model. This is not inherent to MCIA method itself, and provides a prominent reminder that no single ML method is superior to all others, and that selection of training parameters plays a crucial role in the overall method performance.

Going forward, we are committed to performing comparable experiments on a yearly basis that will serve as testing sets for future prediction contests. We believe that this collaborative and innovative approach will create a hub for immunologists to push for novel models of immunity against Tdap boost. We expect the resultant models will also be relevant for other vaccinology studies. Contestants from the research community that are interested in participating are encouraged to contact us via and check the website (www.cmi-pb.org) for the upcoming contest information.

## Methods

### Study design and human participants

Human volunteers that were primed with either the aP or wP vaccination during childhood were recruited. All participants provided written informed consent before donation and were eligible for Tdap (aP) booster vaccination containing tetanus toxoid (TT), diphtheria toxoid (DT), and acellular Pertussis that contains inactivated pertussis toxin (PT) and cell surface proteins of Bordetella pertussis including filamentous hemagglutinin (FHA), fimbriae 2/3 (Fim2/3), pertactin (PRN). Longitudinal blood samples were collected pre-booster vaccination (day 0) and post-booster vaccination after 1, 3, 7, and 14 days.

### PBMC and plasma extraction

Whole blood samples (with heparin) were centrifuged at 1850 rpm for 15 min with breaks off. Subsequently, the upper fraction (plasma) was collected and stored at -80°C. PBMCs were isolated by density gradient centrifugation using Ficoll-Paque PLUS (Amersham Biosciences). 35 mL of RPMI 1640 medium (RPMI, Omega Scientific) diluted blood was slowly layered on top of 15 mL Ficoll-Paque PLUS. Samples were spinned at 1850 rpm for 25 min with breaks off. Then, PBMC layers were aspirated and two PBMC layers per donor were combined in a new tube together with RPMI. Samples were spinned at 1850 rpm for 10 min with a low break. Cell pellets of the same donors were combined and washed with RPMI and spinned at 1850 rpm for 10 min with breaks off. Finally, PBMCs were counted using trypan blue and a hemocytometer and, after another spin, resuspended in FBS (Gemini) containing 10% DMSO (Sigma-Aldrich) and stored in Mr. Frosty cell freezing container overnight at -80°C. The next day, samples were stored at liquid nitrogen until further use.

### Plasma antibody measurements

Pertussis antigen-specific antibody responses were quantified in human plasma by performing an indirect serological assay with xMAP Microspheres (details described in xMAP Cookbook, Luminex 5^th^ edition). Pertussis, Tetanus, and Diphtheria antigens (PT, PRN, Fim2/3, TT, and DT (all from List Biological Laboratories) and FHA (Sigma) and as a negative control Ovalbumin (InvivoGen) were coupled to uniquely coded beads (xMAP MagPlex Microspheres, Luminex Corporation). PT was inactivated by incubation with 1% formaldehyde (PFA) at 4°C for 1 h. 1% PFA PT and TT were then purified using Zeba spin desalting columns (ThermoFisher). The antigens were coupled with each unique conjugated microsphere using the xMAP Antibody Coupling Kit (Luminex Corporation). Plasma was mixed with a mixture of each conjugated microsphere, and WHO International Standard Human Pertussis antiserum was used as a reference standard (NIBSC, 06/140). Subsequently, the mixtures were washed with 0.05% TWEEN20 in PBS (Sigma-Aldrich) to exclude non-specific antibodies, and targeted antibodies responses were detected via anti-human IgG-PE, IgG1-PE, IgG2-PE, IgG3-PE, IgG4-PE (all from SouthernBiotech) and human IgE-PE (ThermoFisher). Antibody details are shown in **Table S4**. Samples were subsequently measured on a FLEXMAP 3D instrument (Luminex Corporation), and the log(10) of the median fluorescence intensity (MFI) was calculated.

### PBMC cell frequencies

Cryopreserved PBMC were thawed by incubating cryovials at 37°C for 1 min and stained with the viability marker Cisplatin. Subsequently, PBMCs were incubated with an antibody mixture for 30 min. After washing, PBMCs were fixed in PBS (Thermo Fisher) with 2% PFA (Sigma-Aldrich) overnight at 4°C. The next day, PBMCs were stained with an intracellular antibody mixture after permeabilization using saponin-based Perm Buffer (eBioscience). After washing, cellular DNA was labeled with Cell-ID Intercalator-Ir (Fluidigm) and cell pellets were resuspended in 1:10 EQ Beads (Fluidigm) in 1 mL MiliQ water. Samples were measured using a Helios mass cytometer (Fluidigm). Antibody details are shown in **Table S4**. Twenty One different PBMC cell subsets were identified using the unsupervised gating approach DAFi^35^ with the exception of antibody-secreting cells (ASCs), which were manually gated as CD45^+^Live^+^CD14^-^CD3^-^CD19^+^CD20^-^CD38^+^ cells. Gating was performed using FlowJo (BD, v10.7).

### Plasma cytokine concentrations

Plasma samples were randomly distributed on 96 well plates for quantification. 276 different proteins (immuno-oncology, immune response, and metabolism Olink panels) were quantified by Analysis Lab at Olink Proteomics. Protein quantification involved the Proximity Extension Assay (PEA) technology^36^. Briefly, the plasma was incubated with oligonucleotides labeled antibodies targeting the proteins of interest. The oligonucleotides of matched oligonucleotides-antibodies-antigen will bind to each other, enabling amplification and thereby quantification by qPCR. Ct values from the qPCR were used to calculate Normalized Protein eXpression (NPX), a relative quantification unit to report protein expression levels in plasma samples.

### RNA sequencing

Per sample, 6 million PBMCs were lysed using QIAzol Lysis Reagent (Qiagen). Samples were stored at -80°C until RNA extraction. RNA was extracted using the miRNeasy Mini Kit (Qiagen) including DNase treatment according to the manufacturer’s instructions. 500 ng of RNA was used for RNA sequencing (RNAseq) library preparation. Library preparation was performed using the TruSeq Stranded mRNA Library Prep Kit (Illumina). Libraries were sequenced on a HiSeq3000 (Illumina) system.

### Bioinformatics RNA sequencing

The pair-end reads that passed Illumina filters were further filtered for reads aligning to tRNA, rRNA, adapter sequences, and spike-in controls. The remaining reads were aligned to the GRCh38 reference genome and Gencode v27 annotations using STAR (v2.6.1)^37^. DUST scores were calculated with PRINSEQ Lite (v0.20.3)^38^, and low-complexity reads (DUST > 4) were removed from the BAM files. The alignment results were parsed via the SAMtools^39^ to generate SAM files. Read counts to each genomic feature were obtained with the featureCounts (v1.6.5)^40^ using the default options along with a minimum quality cut-off (Phred >10).

### Model development

#### Establishing baseline prediction models from the systems vaccinology literature

For literature models that did not present a quantification of the gene sets, a gene set output score was developed. The first step of the calculation was to separate genes that were up- and down-regulated. Next, for each specimen, *i* the TPM normalized gene expression counts were summed for the upregulated genes (*Sum*_*UP*_) and the downregulated genes (*Sum*_*DOWN*_). Then the difference between **Sum*_UP_* and *Sum_DOWN_* was calculated for each specimen:

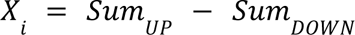

The average (*Aug*) and the standard deviation (*Std*) of the TPM normalized gene expression was calculated across all specimens as well as the square root of the total number of specimens *N*. Finally, a standard score (*zscore*) was calculated for each specimen:

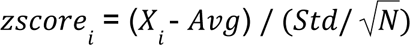

If there were only upregulated genes, or if it could not be determined whether the genes in the gene signature were up- or down-regulated, the sum of the genes in the signature was simply used for the calculation of the *zscore*.

#### Establishing purpose-built models using JIVE

To develop models using JIVE we used harmonized datasets for transcriptome, antibody levels and cytokine levels and located sample values for every variable, in other words, complete datasets which resulted in 13 individuals (**Figure 2C**). Decomposition with JIVE was then applied by using the r.jive package with *jive(omics, rankJ=10, rankA = rep(10, 3)), method = “given”, conv = “default”, maxiter = 100, showProgress=FALSE)* which resulted in 10 factors and saved the factor loading values^41^. In order to make a model from a reduced representation of the omics datasets, each omic was multiplied by its corresponding factor loadings to generate factor scores. During training, these factor scores were used as input into various models and approaches from scikit-learn python package^42^, specifically, we used basic linear regression (LinearRegression), lasso (Lasso), elastic net (ElasticNet), and lastly, lasso and elastic net with automatic hyperparameter tuning via cross-validation (LassoCV and ElasticNetCV). For methods using automatic hyperparameter tuning, candidate values were not specified, therefore using the internal heuristics, which tests hyperparameter values of various magnitudes. During the prediction steps, the testing data is projected onto the lower dimensional space by multiplying the factor loadings by the omics datasets. These new factor scores were then used as input for predictions and finally ranked before contest submission.

#### Establishing purpose-built models using MCIA

The MICE (Multiple Imputation by Chained Equations) method was employed to replace missing data values in the harmonized dataset (**Figure 2D**). Specifically, transcriptome data was utilized to impute missing values in other data modalities through the application of MICE. We utlilized MICE imputed data to construct models. We implemented MCIA using the mbpca function in the *mogsa* package^29,43^. We generated 10 low-dimension multi-omics factor scores for training datasets. Each multi-omics factor was derived through a linear combination of the original features (e.g., genes or proteins) extracted from the input data. Subsequently, global scores were computed for the test dataset, capturing the overall representation or summary of the data in relation to the underlying factors identified from the training data. This was accomplished by utilizing factor loadings from the training dataset and feature scores from the test dataset. For each task, we constructed a prediction model utilizing a general linear model with lasso regularization using the *glmnet* library^44^. We used the feature scores as input data and the prediction task values as response variables, generating separate predictive models for each task.

## Data availability

The training and test datasets used for the first challenge can be accessible through a Zenodo repository at https://doi.org/10.5281/zenodo.7702790. The repository includes detailed information on the datasets, challenge tasks, submission format, descriptions, and access to the necessary data files that contestants used to develop their predictive models and make predictions.

## Code availability

The codebases generated for the first CMI-PB challenge can be accessed through our GitHub group (https://github.com/CMI-PB). The codebase for normalizing antibody titer data is available at https://github.com/CMI-PB/2021-Ab-titer-data-normalisation, while the code for standardizing data and generating computable matrices is available at https://github.com/CMI-PB/cmi-pb-multiomics.

Additionally, the codes for all models submitted for the first CMI-PB challenge are available, including those identified from the literature. All 24 models derived using the literature-based survey are available at https://github.com/CMI-PB/literature_models_first_challenge. The codebase for the JIVE models is available at https://github.com/CMI-PB/cmi-pb-multiomics and the codebase for the MCIA-based models can be found at https://github.com/CMI-PB/mcia-model.

## Acknowledgments

We are grateful to the La Jolla Institute for Immunology Flow Cytometry and Bioinformatics core facilities for their services. The authors would like to thank all donors who participated in the study and the clinical studies group staff - particularly Gina Levi - for all the invaluable help. Research reported in this publication was supported by the National Institute of Allergy and Infectious Diseases of NIH under award no. U01AI150753, U01AI141995 and U19AI142742.

## Author’s contributions

**Study Conception and Design:** BP

**Formal analysis and Interpretation of Results:** PS, JR, AK, MR, BG, LG, FA, BP

**Data Collection and Experimentation:** FSC, LW, MA, RdSA

**Database and Infrastructure:** PS, JO, BH, JaG

**Supervision:** LG, BG, LRO, AS, FA, SK, BP

**Ontology and Curation:** FSC, HGO, JO

**Internal review of manuscript:** SO, RN, MK

**All authors wrote, edited, and reviewed the manuscript.**

## Notes

### Competing Interest Statement

The authors have declared no competing interest.

## References

1. Aghaeepour, N. et al. Critical assessment of automated flow cytometry data analysis techniques. Nat. Methods 10, 228–238 (2013).

2. Ioannidis, J. P., Ntzani, E. E., Trikalinos, T. A. & Contopoulos-Ioannidis, D. G. Replication validity of genetic association studies. Nat. Genet. 29, 306–309 (2001).

3. Eckstein, M. K. et al. The interpretation of computational model parameters depends on the context. eLife 11, e75474 (2022).

4. Shibley, G. S. & Hoelscher, H. Studies on whooping cough : I. Type-specific (s) and dissociation (r) forms of hemophilus pertussis. J. Exp. Med. 60, 403–418 (1934).

5. Edwards, K. M. Challenges to Pertussis Control. Pediatrics 144, (2019).

6. Domenech de Cellès, M., Magpantay, F. M. G., King, A. A. & Rohani, P. The impact of past vaccination coverage and immunity on pertussis resurgence. Sci Transl Med 10, (2018).

7. Klein, N. P., Bartlett, J., Fireman, B. & Baxter, R. Waning Tdap Effectiveness in Adolescents. Pediatrics 137, e20153326 (2016).

8. Klein, N. P., Bartlett, J., Rowhani-Rahbar, A., Fireman, B. & Baxter, R. Waning protection after fifth dose of acellular pertussis vaccine in children. N Engl J Med 367, 1012–9 (2012).

9. Witt, M. A., Arias, L., Katz, P. H., Truong, E. T. & Witt, D. J. Reduced risk of pertussis among persons ever vaccinated with whole cell pertussis vaccine compared to recipients of acellular pertussis vaccines in a large US cohort. Clin Infect Dis 56, 1248–54 (2013).

10. Wilk, M. M. et al. Immunization with whole cell but not acellular pertussis vaccines primes CD4 T. Emerg Microbes Infect 8, 169–185 (2019).

11. Ross, P. J. et al. Relative contribution of Th1 and Th17 cells in adaptive immunity to Bordetella pertussis: towards the rational design of an improved acellular pertussis vaccine. PLoS Pathog 9, e1003264 (2013).

12. da Silva Antunes, R., et al. Th1/Th17 polarization persists following whole-cell pertussis vaccination despite repeated acellular boosters. J Clin Invest 128, 3853–3865 (2018).

13. da Silva Antunes, R., et al. Development and Validation of a Bordetella pertussis Whole-Genome Screening Strategy. J Immunol Res 2020, 8202067 (2020).

14. da Silva Antunes, R., et al. >A system-view of Bordetella pertussis booster vaccine responses in adults primed with whole-cell versus acellular vaccine in infancy. JCI Insight 6, (2021).

15. Gayen, S., Maclary, E., Hinten, M. & Kalantry, S. Sex-specific silencing of X-linked genes by Xist RNA. Proc. Natl. Acad. Sci. U. S. A. 113, E309–318 (2016).

16. Fourati, S. et al. Pan-vaccine analysis reveals innate immune endotypes predictive of antibody responses to vaccination. Nat Immunol 23, 1777–1787 (2022).

17. Tsang, J. S. et al. Global analyses of human immune variation reveal baseline predictors of postvaccination responses. Cell 157, 499–513 (2014).

18. Kotliarov, Y. et al. Broad immune activation underlies shared set point signatures for vaccine responsiveness in healthy individuals and disease activity in patients with lupus. Nat Med 26, 618–629 (2020).

19. Fourati, S. et al. Pre-vaccination inflammation and B-cell signalling predict age-related hyporesponse to hepatitis B vaccination. Nat Commun 7, 10369 (2016).

20. Furman, D. et al. Apoptosis and other immune biomarkers predict influenza vaccine responsiveness. Mol Syst Biol 9, 659 (2013).

21. HIPC-CHI Signatures Project Team & HIPC-I Consortium. Multicohort analysis reveals baseline transcriptional predictors of influenza vaccination responses. Sci. Immunol. 2, eaal4656 (2017).

22. di Iulio, J., Bartha, I., Spreafico, R., Virgin, H. W. & Telenti, A. Transfer transcriptomic signatures for infectious diseases. Proc Natl Acad Sci U A 118, (2021).

23. Bartholomeus, E. et al. Transcriptome profiling in blood before and after hepatitis B vaccination shows significant differences in gene expression between responders and non-responders. Vaccine 36, 6282–6289 (2018).

24. Qiu, S. et al. Significant transcriptome and cytokine changes in hepatitis B vaccine non-responders revealed by genome-wide comparative analysis. Hum Vaccin Immunother 14, 1763–1772 (2018).

25. Franco, L. M. et al. Integrative genomic analysis of the human immune response to influenza vaccination. Elife 2, e00299 (2013).

26. Zhang, L. et al. Cross-platform comparison of immune-related gene expression to assess intratumor immune responses following cancer immunotherapy. J Immunol Methods 494, 113041 (2021).

27. Lock, E. F., Hoadley, K. A., Marron, J. S. & Nobel, A. B. Joint and individual variation explained (jive) for integrated analysis of multiple data types. Ann Appl Stat 7, 523–542 (2013).

28. White, I. R., Royston, P. & Wood, A. M. Multiple imputation using chained equations: Issues and guidance for practice. Stat Med 30, 377–99 (2011).

29. Meng, C., Kuster, B., Culhane, A. C. & Gholami, A. M. A multivariate approach to the integration of multi-omics datasets. BMC Bioinformatics 15, 162 (2014).

30. Meng, C. et al. Dimension reduction techniques for the integrative analysis of multi-omics data. Brief Bioinform 17, 628–41 (2016).

31. Meyer, P. & Saez-Rodriguez, J. Advances in systems biology modeling: 10 years of crowdsourcing DREAM challenges. Cell Syst. 12, 636–653 (2021).

32. Moult, J., Fidelis, K., Rost, B., Hubbard, T. & Tramontano, A. Critical assessment of methods of protein structure prediction (CASP)--round 6. Proteins 61 Suppl 7, 3–7 (2005).

33. Aiello, A. et al. Immunosenescence and Its Hallmarks: How to Oppose Aging Strategically? A Review of Potential Options for Therapeutic Intervention. Front. Immunol. 10, 2247 (2019).

34. Verschoor, C. P. et al. Advanced biological age is associated with improved antibody responses in older high-dose influenza vaccine recipients over four consecutive seasons. Immun. Ageing A 19, 39 (2022).

35. Lee, A. J. et al. DAFi: A directed recursive data filtering and clustering approach for improving and interpreting data clustering identification of cell populations from polychromatic flow cytometry data. Cytom. Part J. Int. Soc. Anal. Cytol. 93, 597–610 (2018).

36. Assarsson, E. et al. Homogenous 96-plex PEA immunoassay exhibiting high sensitivity, specificity, and excellent scalability. PloS One 9, e95192 (2014).

37. Dobin, A. et al. STAR: ultrafast universal RNA-seq aligner. Bioinforma. Oxf. Engl. 29, 15–21 (2013).

38. Schmieder, R. & Edwards, R. Quality control and preprocessing of metagenomic datasets. Bioinforma. Oxf. Engl. 27, 863–864 (2011).

39. Li, H. et al. The Sequence Alignment/Map format and SAMtools. Bioinforma. Oxf. Engl. 25, 2078–2079 (2009).

40. Liao, Y., Smyth, G. K. & Shi, W. featureCounts: an efficient general purpose program for assigning sequence reads to genomic features. Bioinforma. Oxf. Engl. 30, 923–930 (2014).

41. O’Connell, M. J. & Lock, E. F. R.JIVE for exploration of multi-source molecular data. Bioinforma. Oxf. Engl. 32, 2877–2879 (2016).

42. Pedregosa, F. et al. Scikit-learn: Machine Learning in Python. J. Mach. Learn. Res. 12, 2825–2830 (2011).

43. Meng, C. et al. MOGSA: Integrative Single Sample Gene-set Analysis of Multiple Omics Data. Mol Cell Proteomics 18, S153–S168 (2019).

44. Friedman, J., Hastie, T. & Tibshirani, R. Regularization Paths for Generalized Linear Models via Coordinate Descent. J. Stat. Softw. 33, 1–22 (2010).

