## Supplementary Materials for "A systems vaccinology resource to develop and test computational models of immunity"

#### INDEX

**Supplementary tables:**

S1. List of previously identified Tdap Immune signatures represented as tasks for 1st challenge.

S2. Implemented prediction methods from literature.

S3. List of antibodies

**Supplementary figures:**

S1. CMI-PB database schema.

S2. A plot of antibody titer data prior to normalization.

S3. A plot of antibody titer data post normalization.

S4. Identification of published methods to predict vaccine responses.

**Reference**

### Supplementary Tables

#### Table S1: List of previously identified Immune signatures represented as tasks for the first challenge.

| | Task Title | Task statement | Signature | Reference |
|---|---|---|---|---|
| **Antibody level tasks** | | | | |
| 1 | IgG-PT_D14 | IgG levels against Pertussis Toxin (PT) antigen in plasma on 14 days post booster vaccination | Vaccine-induced immunity for B. pertussis | [1]] |
| 2 | IgG-FHA_D14 | IgG levels against Filamentous hemagglutinin (FHA) antigen in plasma on 14 days post booster vaccination | Vaccine-induced immunity for *B. pertussis* | [1]] |
| 3 | IgG-Pertactin_D14 | IgG levels against Pertactin antigen in plasma on 14 days post booster vaccination | Vaccine-induced immunity for *B. pertussis* | Figure 5 in [1] |
| 4 | IgG1-PT_D14 | IgG1 levels against PT antigen in plasma on 14 days post booster vaccination | Vaccine-induced immunity for *B. pertussis* | Figure 5 in [1] |
| 5 | IgG1-FHA_D14 | IgG1 levels against FHA antigen in plasma on 14 days post booster vaccination | Vaccine-induced immunity for *B. pertussis* | Figure 5 in [1] |
| 6 | IgG4-PT_D14 | IgG4 levels against PT antigen in plasma on 14 days post booster vaccination | Acellular Pertussis (aP) vs. whole-cell Pertussis (wP) response | Figure 5 in [1] |
| 7 | IgG4-FHA_D14 | IgG4 levels against FHA antigen in plasma on 14 days post booster vaccination | aP vs. wP response | Figure 5 in [1] |
| **Cell frequency tasks** | | | | |
| 8 | Plasmablast_D7 | Plasmablast cells on day 7 post-booster vaccination | Vaccine-induced immunity for *B. pertussis* | Figure 3 in [1] |
| 9 | CD4TCM_D3 | CD4 TCM cells on 3 days post booster vaccination | Vaccine-induced immunity for *B. pertussis* | Figure 3 in [1] |
| 10 | Monocytes_D3 | Monocytes on day 1 post-booster vaccination | Vaccine-induced immunity for *B. pertussis* | Figure 3 in [1] |
| **Gene expression tasks** | | | | |
| 11 | CCL3_D3 | CCL3 on day 3 post booster vaccination | aP vs. wP response | Figure 7 in [1] |
| 12 | IL6_D3 | IL6 on day 3 post-booster vaccination | aP vs. wP response | Figure 7 in [1] |
| 13 | NFKBIA_D7 | NFKBIA at day 7 post booster vaccination | aP vs. wP response | Figures 6 and 8 in [1] |
| 14 | XIST_D14 | XIST on day 14 post-booster vaccination | Biological sex-specific marker | [2] |

**Table S2: Implemented prediction methods for the first challenge.** Models 1-24: Supervised methods based on published vaccine response prediction studies. Models 25-43: unsupervised methods based on Multi-omics dimension reduction.

| | Model title | Short Description | Method type | Output score | Experimental data | Reference |
|---|---|---|---|---|---|---|
| 1 | avey_2017_gene_sig | 9-gene signature (RAB24, GRB2, DPP3,ACTB, MVP, DPP7, ARPC4, PLEKHB2, and ARRB1) | Classification (Diagonal Linear Discriminant Analysis: DLDA) | geneset signature score | **Experimental data:** PBMC and whole blood gene expression, cell subset frequencies, antibody titers<br><br>**Pathogen:** Influenza virus | Avey et al. 2017 [3] |
| 2 | avey_2017_M54 | B-cell signaling Blood transcriptome module (BTM; Module M54) | Classification (DLDA) | geneset signature score | | |
| 3 | avey_2017_M42 | Platelet activation (III) (Module M42) | Classification (DLDA) | geneset signature score | | |
| 4 | avey_2017_M33 | Inflammatory response Module (M33) | Classification (DLDA) | geneset signature score | | |
| 5 | kotliarov_2020_TGSig | 10 gene signature (C2orf63, CD101, ENPP1, RETN, SMC1A, ADAM12, EPHB1, PAPSS2, LONP2, C15orf57) | gene signature score | geneset signature score | **Experimental data:** PBMC and whole blood gene expression, cell subset frequencies, cell surface proteins, antibody titers<br><br>**Pathogen:** Influenza virus, Yellow fever virus | Kotliarov et al. 2020 [4] |
| 6 | kotliarov_2020_SLE-Sig | Systemic lupus erythematosus (SLE-Sig) gene signature | gene signature score | geneset signature score | | |
| 7 | kotliarov_2020_IFN-I-DCact | IFN-I-DCact gene signature | gene signature score | geneset signature score | | |
| 8 | tsang_2014_DLDA_top2 | DLDA model (2 cell populations | Classification (DLDA) | Class probabilities | **Experimental data:** PBMC gene expression, cell subset frequencies, antibody titers<br><br>**Pathogen:** Influenza virus | Tsang et al. 2014 [5][24725414] |
| 9 | tsang_2014_DLDA_top5 | DLDA model (5 cell populations) | Classification (DLDA) | Class probabilities | | |
| 10 | fourati_2015_BioAge | BioAge | Classification (Naïve Bayes) | Class probabilities | **Experimental data:** Whole blood gene expression, cell subset frequencies, serum proteins, antibody titers<br><br>**Pathogen:** Hepatitis B virus | Fourati et al. 2015 [6] |
| 11 | fourati_2015_M1 | M1 of BioAge | Classification (Naïve Bayes) | Class probabilities | | |
| 12 | fourati_2015_M1+M16 | M1+M16 of BioAge | Classification (Naïve Bayes) | Class probabilities | | |

| | | | | | | |
|---|---|---|---|---|---|---|
| 13 | fourati_2015_NB | Naïve Bayes classifier (15 DEGs) | Classification (Naïve Bayes) | The ratio between the post-probabilities and the log odds in the logistic model | | |
| 14 | fourati_2015_LR | Logistic regression (4 cell populations) | Regression (logistic) | The ratio between the post-probabilities and the log odds in the logistic model | | |
| 15 | furman_2013_age | gene signature scores Age | Regression (Elastic net) | age | **Experimental data:** Whole blood gene expression, cell subset frequencies serum cytokines, antibody titers, hemagglutinin peptides<br><br>**Pathogen:** Influenza virus | Furman et al. 2013 [7] |
| 16 | iulio_2021_HBV_transfer_sig | HBV pre-vaccine transfer signature | Classification (Random Forest) | geneset signature score | **Experimental data:** PBMC and whole blood gene expression<br><br>**Pathogen:** Influenza virus, Hepatitis B virus, Mycobacterium tuberculosis | Iulio et al. 2021 [8] |
| 17 | iulio_2021_Inf_M_transfer_sig | Influenza M pre-vaccine transfer signature | Classification (Random Forest) | geneset signature score | | |
| 18 | iulio_2021_Inf_F_transfer_sig | Influenza F pre-vaccine transfer signature | Classification (Random Forest) | geneset signature score | | |
| 19 | iulio_2021_TB_transfer_sig | TB pre-vaccine transfer signature | Classification (Random Forest) | geneset signature score | | |
| 20 | fourati_2021_RF | Random Forest model (top 500 varying genes) | Classification (Random Forest) | Class probabilities | **Experimental data:** PBMC and whole blood gene expression, cell surface proteins, cell subset frequencies, antibody titers<br><br>**Pathogen:** Influenza, smallpox, Yellow fever virus, Pneumococcal meningococcal | Fourati et al. 2021 [9] |
| 21 | bartholomeus_2018_gene_sig | 23 differentially expressed genes | Classification (Random Forest) | Class probabilities | **Experimental data:** Whole blood gene expression, absolute numbers of white blood cells, red blood cells, and platelets, antibody titers<br><br>**Pathogen:** Hepatitis B virus | Bartholomeus et al. 2018 [10][30205979] |
| 22 | bartholomeus_2018_NB | Naïve Bayes classifier (first 5 PCs) | Classification (Naïve Bayes) | Class probabilities | | |
| 23 | qui_2018_gene_sig | 55 up- and 15 down-regulated genes | gene signature score | geneset signature score | **Experimental data:** PBMC gene expression, antibody titers<br><br>**Pathogen:** Hepatitis B virus | Qui et al. 2018 [11] |
| 24 | franco_2013_gene_sig | 49 genes correlated | gene signature score | geneset signature score | **Experimental data:** Whole blood gene expression, | Franco et |

| | | | | | | |
|---|---|---|---|---|---|---|
| | | with antibody response | | | whole-genome genotyping, antibody titers<br><br>**Pathogen:** Influenza virus | al. 2013 [12] |
| 25 | jive.elastic_net_cv | Elastic net regression with cross-validati on | Regression (Elastic net) | | **Experimental data:** PBMC gene expression, cell frequencies, antibody titers | Loc k et al. 201 3 [13] |
| 26 | jive.elastic_net | Elastic net regression | Regression (Elastic net) | | | |
| 27 | jive.lasso_cv | lasso regression with cross-validati on | Regression (lasso) | | | |
| 28 | jive.lasso | lasso regression | Regression (lasso) | | | |
| 29 | jive.lr | Linear regression | Regression (linear) | | | |
| 30 | baseline | | Regression (lasso) | | Baseline tasks, clinical and demographic features | |
| 31 | MCIAbasic | | Regression (lasso) | | **Experimental data:** transcriptome, cell frequencies, antibody levels, cytokines | |
| 32 | MCIAplus | | Regression (lasso) | | Factors from the MCIAbasic model and features from the baseline model | |

**Table S3**. Antibodies details

| Target | Conjugate | Host | Target | Clone | Catalog | Vendor | Application | Dilution |
|---|---|---|---|---|---|---|---|---|
| IgG | PE | Mouse | Human | JDC-10 | 9040-09 | Southern Biotech | Antibody response | 1/50 |
| IgG1 | PE | Mouse | Human | HP6001 | 9054-09 | Southern Biotech | Antibody response | 1/50 |
| IgG2 | PE | Mouse | Human | HP6025 | 9070-09 | Southern Biotech | Antibody response | 1/20 |
| IgG3 | PE | Mouse | Human | HP6050 | 9210-09 | Southern Biotech | Antibody response | 1/10 |
| IgG4 | PE | Mouse | Human | HP6025 | 9200-09 | Southern Biotech | Antibody response | 1/10 |
| IgE | PE | Mouse | Human | BE5 | MA1-10 375 | Thermo Fisher | Antibody response | 1/10 |
| CD45 | 89Y | | Human | HI30 | 308900 3B | Fluidigm | CyTOF | |
| CD3 | 115In | | Human | UCHT1 | 300443 | Biolegend | CyTOF | |
| CD19 | 142Nd | | Human | HIB19 | 314200 1B | Fluidigm | CyTOF | |
| CD38 | 144Nd | | Human | HIT2 | 314401 4B | Fluidigm | CyTOF | |
| CD4 | 145Nd | | Human | RPA-T4 | 314500 1B | Fluidigm | CyTOF | |
| CD20 | 145Nd | | Human | 2H7 | 302302 | Biolegend | CyTOF | |
| CD12 3 | 151Eu | | Human | 6H6 | 315100 1B | Fluidigm | CyTOF | |
| CD45 RA | 155Gd | | Human | HI100 | 315501 1B | Fluidigm | CyTOF | |
| CD1c | 160Gd | | Human | L161 | 331502 | Biolegend | CyTOF | |
| CD33 | 163Dy | | Human | WM53 | 316302 3B | Fluidigm | CyTOF | |

| | | | | | | | |
|---|---|---|---|---|---|---|---|
| CCR7 | 167Er | | Human | G043H7 | 3167009A | Fluidigm | CyTOF |
| CD25 | 169Tm | | Human | M-A251 | 3169003B | Fluidigm | CyTOF |
| CD8a | 172Yb | | Human | RPA-T8 | 301002 | Biolegend | CyTOF |
| CD14 | 173Yb | | Human | 61D3 | 14-0149-82 | Thermo Fisher | CyTOF |
| HLA-DR | 174Yb | | Human | L243 | 3174001B | Fluidigm | CyTOF |
| CD56 | 176Yb | | Human | CMSSB | 3176003B | Fluidigm | CyTOF |
| CD16 | 209Bi | | Human | 3G8 | 3209002B | Fluidigm | CyTOF |

#### Supplementary Figures

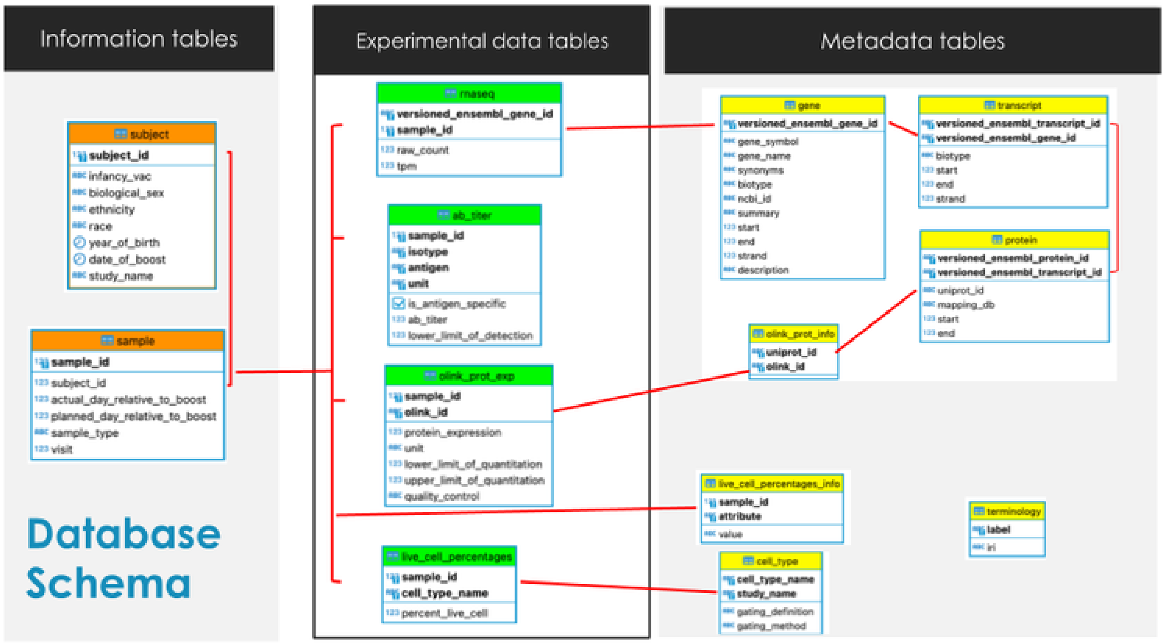

**Figure S1: CMI-PB central database schema.** The database schema is divided into information, experimental data, and metadata tables. The information tables capture subject and sample information, while the experimental data tables capture experimental data for each omics. In the *rnaseq* table, each row represents an Ensembl gene identifier (versioned_ensembl_gene_id) along with specimen id (specimen_id) and values for measured analytes (raw_count and tpm_count). The *ab_titer* table contains a row for each specimen (specimen_id), with the names and values for measured analytes (isotype, is_antigen_specific, antigen, ab_titer, unit, and lower_limit_of_detection) for the antibody titer experiment. The *olink* table contains a row for each specimen (specimen_id), with the names and values for measured analytes (olink_id and protein_expression). Each row in the *live_cell_percentages* table represents a specimen (specimen_id) along with the names and values for measured analytes (cell_type_name and percent_live_cell). Information on the mapping between olink_id and uniprot_id can be extracted using the *olink_prot_info* table. The metadata tables capture information about ID mapping between CMI-PB data and external databases. The gene, transcript, and protein tables map Ensembl gene, transcript, and protein IDs. These Ensembl IDs are then mapped to UniProt IDs. The *live_cell_percentages_info* and *cell_type* tables provide information about gating information and the experimental technique used to run cell frequency experiments.

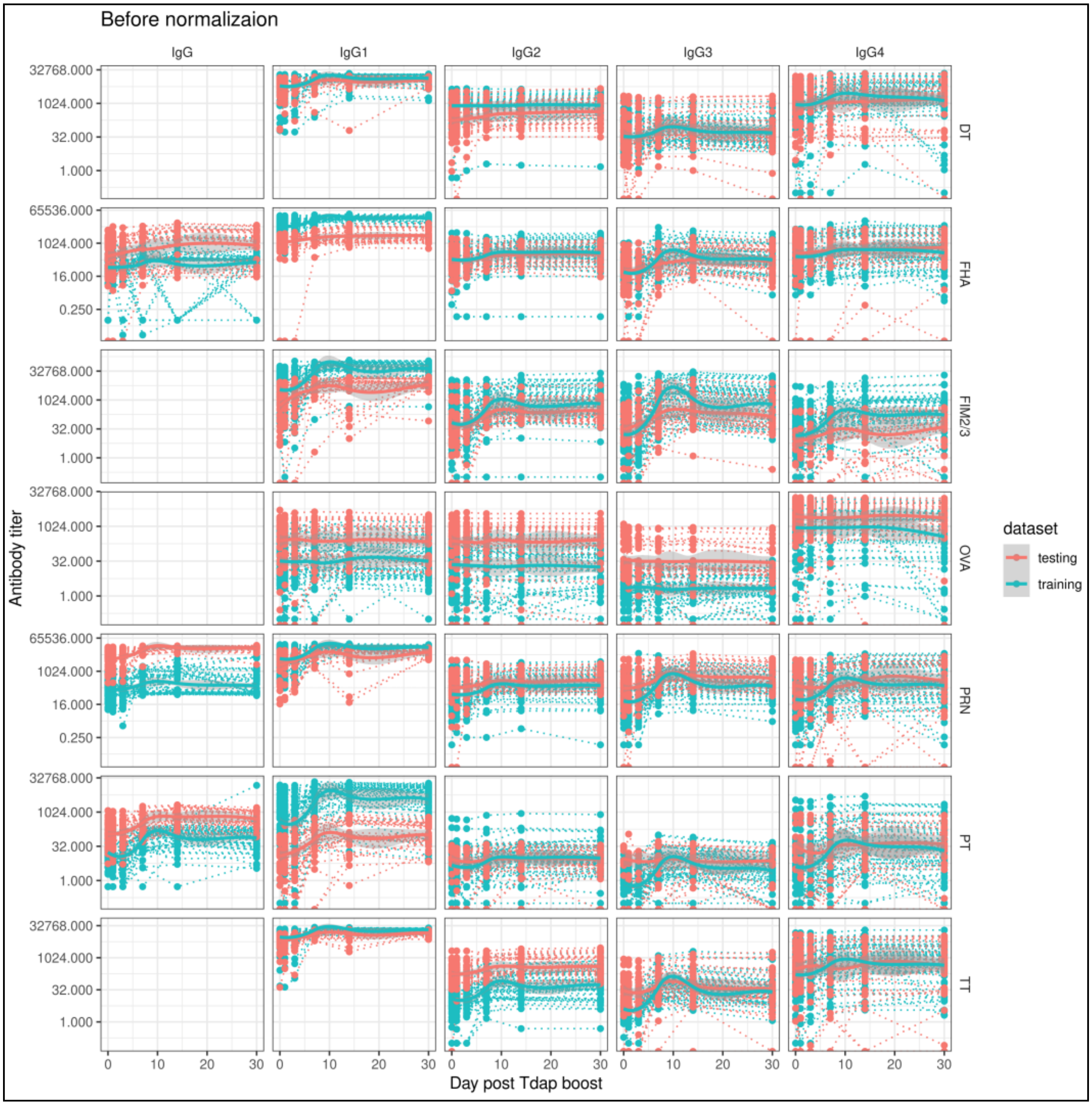

**Figure S2: Plot of antibody titer data prior to normalization.** Individual plots show the pre- and post-immune response of IgG and its subtypes against antigens. The average titer values differed between the train and test datasets.

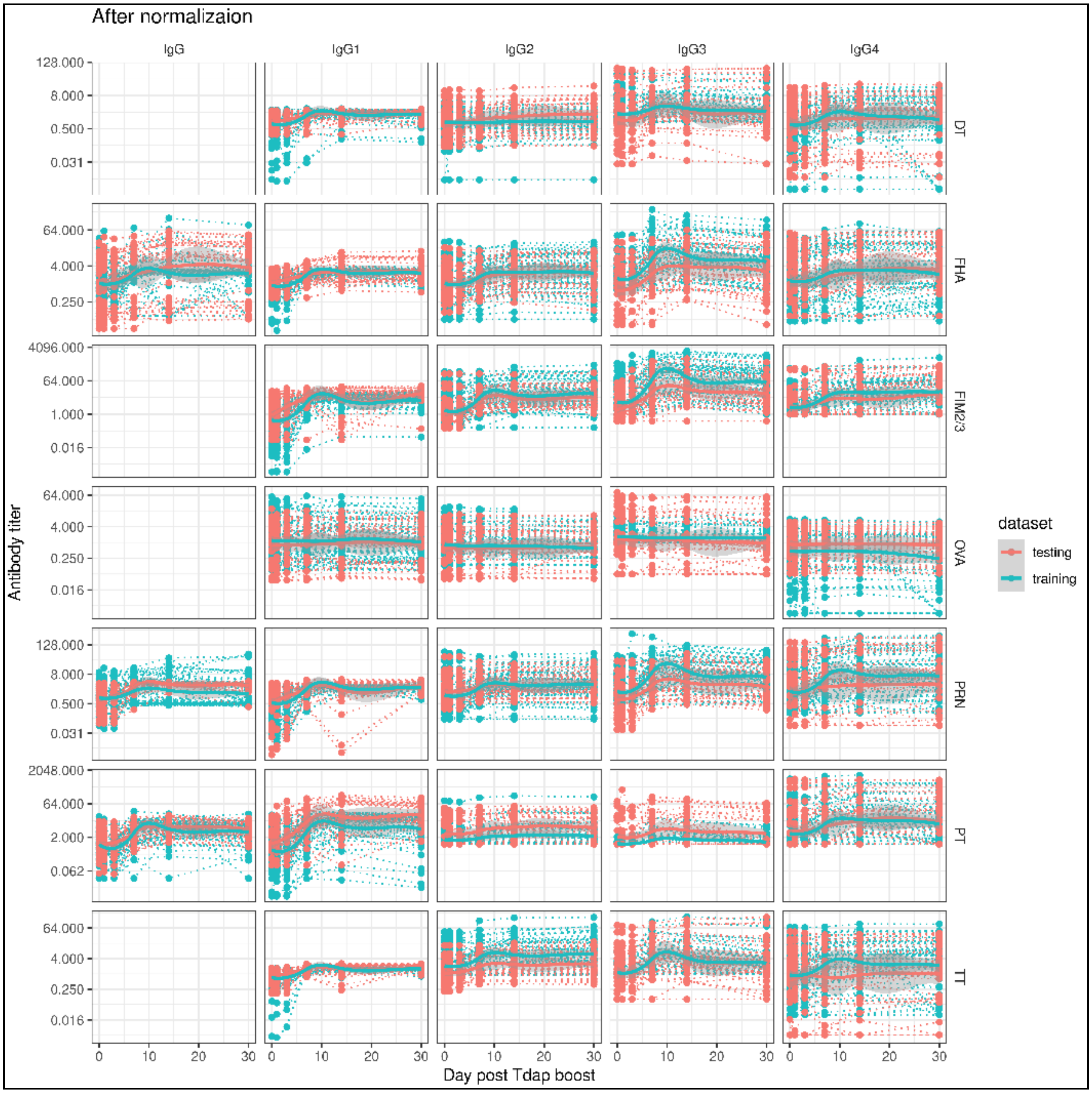

**Figure S3: Plot of antibody titer data after normalization.** We performed separate standardization of the antibody data in the train and test datasets, using the baseline median as the normalization factor. This approach allowed more direct comparisons of the normalized datasets between the train and test datasets.

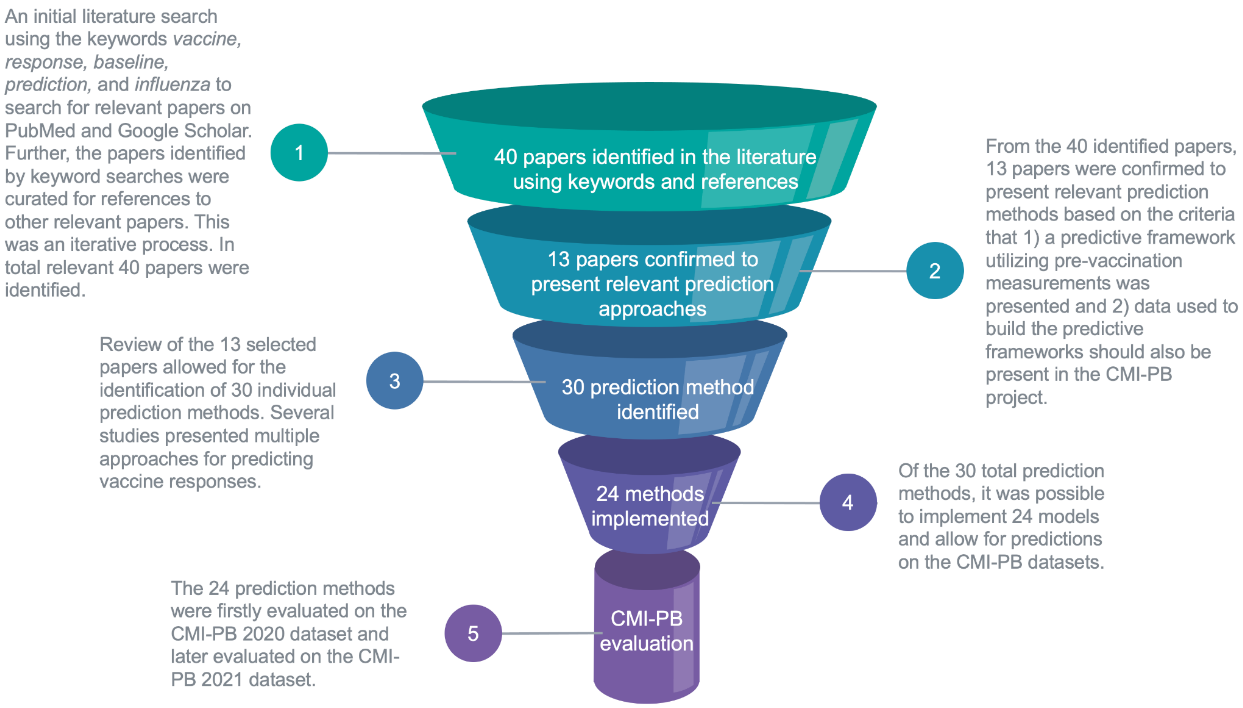

**Figure S4: Identification of published methods to predict vaccine responses.** The literature search identified previously presented methods for predicting vaccine responses. First, 40 papers were selected from the literature search using keyword searches and relevant references in other papers. From the 40 papers, 13 studies were confirmed to present relevant prediction methods based on a literature search for published vaccine response prediction methods. For several of the 13 relevant studies, multiple prediction methods were presented. In total, 24 prediction methods were implemented from 10 relevant 13 studies (mentioned in **Table S1**). The prediction methods that obtained significant results for both performance metrics in the CMI-PB train datasets were also evaluated in the CMI-PB test dataset used for the first challenge.

**References:**

1. da Silva Antunes, R. *et al.* A system-view of Bordetella pertussis booster vaccine responses in adults primed with whole-cell versus acellular vaccine in infancy. *JCI Insight* **6**, (2021).
2. Gayen, S., Maclary, E., Hinten, M. & Kalantry, S. Sex-specific silencing of X-linked genes by Xist RNA. *Proc. Natl. Acad. Sci. U. S. A.* **113**, E309-318 (2016).
3. HIPC-CHI Signatures Project Team & HIPC-I Consortium. Multicohort analysis reveals baseline transcriptional predictors of influenza vaccination responses. *Sci. Immunol.* **2**, eaal4656 (2017).
4. Kotliarov, Y. *et al.* Broad immune activation underlies shared set point signatures for vaccine responsiveness in healthy individuals and disease activity in patients with lupus. *Nat Med* **26**, 618–629 (2020).
5. Tsang, J. S. *et al.* Global analyses of human immune variation reveal baseline predictors of postvaccination responses. *Cell* **157**, 499–513 (2014).
6. Fourati, S. *et al.* Pre-vaccination inflammation and B-cell signalling predict age-related hyporesponse to hepatitis B vaccination. *Nat Commun* **7**, 10369 (2016).
7. Furman, D. *et al.* Apoptosis and other immune biomarkers predict influenza vaccine responsiveness. *Mol Syst Biol* **9**, 659 (2013).
8. di Iulio, J., Bartha, I., Spreafico, R., Virgin, H. W. & Telenti, A. Transfer transcriptomic signatures for infectious diseases. *Proc Natl Acad Sci U A* **118**, (2021).
9. Fourati, S. *et al.* Pan-vaccine analysis reveals innate immune endotypes predictive of antibody responses to vaccination. *Nat Immunol* **23**, 1777–1787 (2022).
10. Bartholomeus, E. *et al.* Transcriptome profiling in blood before and after hepatitis B vaccination shows significant differences in gene expression between responders and non-responders. *Vaccine* **36**, 6282–6289 (2018).
11. Qiu, S. *et al.* Significant transcriptome and cytokine changes in hepatitis B vaccine non-responders revealed by genome-wide comparative analysis. *Hum Vaccin Immunother*

**14**, 1763–1772 (2018).

12. Franco, L. M. *et al.* Integrative genomic analysis of the human immune response to influenza vaccination. *Elife* **2**, e00299 (2013).
13. Lock, E. F., Hoadley, K. A., Marron, J. S. & Nobel, A. B. JOINT AND INDIVIDUAL VARIATION EXPLAINED (JIVE) FOR INTEGRATED ANALYSIS OF MULTIPLE DATA TYPES. *Ann Appl Stat* **7**, 523–542 (2013).
